# The structural extraction of Chinese medical narratives

**DOI:** 10.1101/380626

**Authors:** Rongzhi Zhang, Haifei Zhang, Zhiyu Yao, Zhengxing Huang

## Abstract

Medical narratives document a vast amount of clinical data. This data has a valuable secondary purpose, as it may be used to optimize health service delivery and improve the quality of medical care. However, medical narratives are typically recorded in an unstructured manner, which complicates the process of extracting the structured information required for optimization. In this paper, we address this problem by applying and comparing two models, a rule-based model and a model based on conditional random fields (CRFs), to a data set of Chinese medical narratives. Among 4626 manually annotated Chinese medical narratives, collected from Shanxi Dayi Hospital in China, the rule-based model achieved 95.87% precision, 69.82% recall, and an F-score of 80.80%, and the CRF-based model realized 95.99% precision, 65.11% recall, and a 77.59% F-score. These experimental results demonstrate the efficacy of both proposed models for structural extraction from Chinese medical narratives.

## Introduction

Since the 1960s, natural language processing (NLP) research has been active in the clinical sector. Although medical NLP research has lagged behind more general NLP applications [1], a great deal of meaningful work has been done. Lack of shared data used to impede medical NLP research, so the efforts of [2–5] to make clinical data available for sharing merit note. There is also a large body of work on NLP for clinical information [6–16], including applications related to respiratory illness, radiology, discharge summaries, electrocardiography, echocardiography, pharmacovigilance, postoperative surgical complications, and pathology. This study focused on medical narrative that documented progress notes and daily rounds for patients with cardiovascular diseases. One well-known NLP project, the Language String Project-Medical Language Processor (LSP-MLP) [17–19], extracts and summarizes symptomologies and medication dosage information [20]. Columbia University has developed Medical Language Extraction and Coding System (MEDLEE), an NLP system that identifies clinical information in narrative reports and converts textual information into structured representations [21]. This study shares the objectives of these systems, but differs from the existing approaches in that the proposed method is specifically applied to medical narratives composed using Chinese characters and language.

Despite its late start, Chinese natural language processing has made great progress in the past two decades [22–24]. Huang et al. proposed a segmentation standard for Chinese natural language processing which serves as a preliminary work on the structural extraction of Chinese medical narratives [25]. A group from Harbin Institute of Technology developed an integrated Chinese language processing platform that achieved good results by executing syntactic and semantic parsing modules [26]. Wang et al. presented a method for symptom name recognition from free-text clinical records based on conditional random fields (CRFs) [27]. Xu et al. developed two independent systems of word segmentation and named entity recognition which were also based on CRFs [28].

As a country, China has a well-established and solid foundation for the information construction of medical institutions. The electronic medical record (EMR) system has been widely applied by medical institutions at all levels. The content of EMRs consists of unstructured narrative text as well as structured coded data[29]. Although EMRs contain abundant data for advanced research, existing techniques for processing unstructured text are too inefficient for practical application. Therefore, the key to data mining EMRs is to fulfill the structural extraction of the medical record data, which is accomplished by processing EMR narrative content into structured text to facilitate effective data mining subsequently. Both proposed models, the rule-based model and the CRF model, have been widely used for English medical language processing [30–32]. Because of the difficulties encountered in word segmentation and other ambiguities, natural language processing is more complex and less understood for documents written in Chinese.

In this paper, we evaluate the execution of rule-based and CRF models on Chinese medical narratives and compare the results. Factors limiting the extraction results were also explored. Specifically, we conducted a detailed analysis of the word segmentation process. The unique contributions of this paper are as follows:

1. Instead of relying a single model, we executed and compared a rule-based model and a CRF model;
2. For the structural extraction of Chinese medical narratives with similar writing habits, we observed that the rule-based model achieved higher recall while the CRF-based model attained higher precision;
3. To our knowledge, there has been little focused study of the progress notes and daily records in Chinese medical narratives that were used in this study.

## Materials and methods

### Data sources and overview

We obtained EMRs from the cooperative hospital, Shanxi Dayi Hospital, located in Taiyuan, China. Each EMR is comprised of patient progress notes and daily records sorted by timestamp. Each EMR describes the physical condition and associated diagnostic information of the patient at the time of observation. If applicable, examination results are also included. Owing to variations in EMR reporting criteria instituted by different hospitals, we suggest tailoring preprocessing to the EMR protocols of the target hospital. However, after targeted preprocessing, the structured extraction methods of the proposed method are universal. Table 1 provides an excerpt from one of the EMRs used in this study, as original transcribed in Chinese and translated to English.

**Table 1.** Excerpt of an electronic medical record (EMR) as originally transcribed in Chinese natural language and translated into English.

|  |  |
| --- | --- |
| Original Chinese language<br>daily progress notes<br>and records | 2015-03-28 16:58 日常病程记录<br>今日查房,患者精神、食欲可,咳嗽、咳痰症状改善。查体:体温 38.4℃,血压 120/66mmHg,神志清楚,双肺呼吸音清,未闻及干湿性啰音,心前区无隆起,心率 66 次/分,律齐,各瓣膜听诊区未闻及杂音,腹软,无压痛、反跳痛,双下肢不肿。胸片未见异常。化验示总胆固醇 3.17 mmol/L,甘油三酯 2.27 mmol/L,补充诊断高甘油三酯血症,嘱低脂饮食,继续冠心病二级预防治疗。<br>*** |
|  | 2015-03-31 09:02 主任医师查房记录<br>今日***副主任医师查房,患者无特殊不适。查体:体温 36℃,血压 114/64mmHg,神志清楚,双肺呼吸音清,未闻及干湿性啰音,心前区无隆起,心率 64 次/分,律齐,各 |
|  | 瓣膜听诊区未闻及杂音，腹软，无压痛、反跳痛，双下肢不肿。心脏彩超示左房扩大，三尖瓣关闭不全（轻度），肺动脉压（PASP=31mmHg）。继续冠心病二级预防及抗感染治疗，患者要求明日出院，嘱规律服药，门诊复查。<br>*** |
| English translation of EMR excerpt | 2015-03-28 16:58 Daily progress note<br>Checking the room today, the patient's spirit, appetite, cough, sputum symptoms improved. Physical examination: body temperature 38.4° C; blood pressure 120/66mmHg; conscious; breath sounds clear in both lungs, dry and wet rales not heard; no bulge in precordial area; heart rate 66 beats/min, regularity; noise unheard in all valve auscultation areas; soft abdomen; no tenderness or rebound tenderness; no swelling of both lower extremities; no abnormal chest radiograph. The test showed total cholesterol 3.17 mmol/L, triglyceride 2.27 mmol/L. Additional diagnosis of hypertriglyceridemia. It is suggested to keep a reasonable diet, and to continue secondary treatment of coronary heart disease.<br>***<br>2015-03-31 09:02 Director doctor daily records<br>***, deputy director of the doctor checks the room today, the patient had no special discomfort. Physical examination: body temperature 36 ° C; blood pressure 114/64mmHg; conscious; breath sounds in both lungs; dry and wet rales not heard; precordial no bulge; heart rate 64 beats/min; noise unheard in all valve auscultation areas; soft abdomen; no tenderness or rebound tenderness; no swelling of both lower extremities. Cardiac echocardiography showed enlargement of the left atrium, tricuspid regurgitation (mildness), and pulmonary artery pressure (PASP = 31 mmHg). Continue secondary prevention and anti-infective treatment of coronary heart disease, the patient requested to leave the hospital tomorrow. Regular medication and outpatient review is suggested.<br>*** |

### Data preprocessing

As previously mentioned, targeted preprocessing is applied to sets of EMRs obtained from individual hospitals to standardize the data for universal structured extraction techniques. An EMR, or clinical narrative, contains multiple progress notes and daily records divided into stages according to the timestamp of each notation. In the Chinese language, clauses are independent and complete when separated by commas and periods. Thus, for each stage, or individually timestamped note, the text was separated into independent clauses based on the punctuation. In the final step of preprocessing, the independent clauses were segmented using Jieba, a Chinese word segmentation tool. Table 2 provides an example of an input clause and the segmentation implemented during preprocessing.

**Table 2.** Example of input clause and annotated output after segmentation in preprocessing.

|  | Clause | Annotated Clause* |
| --- | --- | --- |
| Chinese | 双肺呼吸音清 | PF#0#DS#1# {'PF#0#': '双肺呼吸音', 'DS#1#': '清'} |
| English | Lung breath sounds clear | PF#0#DS#1# {'PF#0#': 'Lung breath', 'DS#1#': 'clear'} |

The tags “PF” and “DS” abbreviate “physiological function” and “description,” respectively.

### The rule-based model

#### The pipeline of the rule-based model

Fig 1 presents the structured extraction process of the rule-based model. First, the clauses of Chinese medical narratives are input as test data. Next, the Jieba segmentation tool separated the input clauses into individual terms. Then, if the terms are found in the concept dictionary, they are replaced by code indicating their associated concept and an ordinal number. The rules are then applied to the encoded segments to extract the valid information from the clause and produce the structural results as the final output.

**Fig 1.**
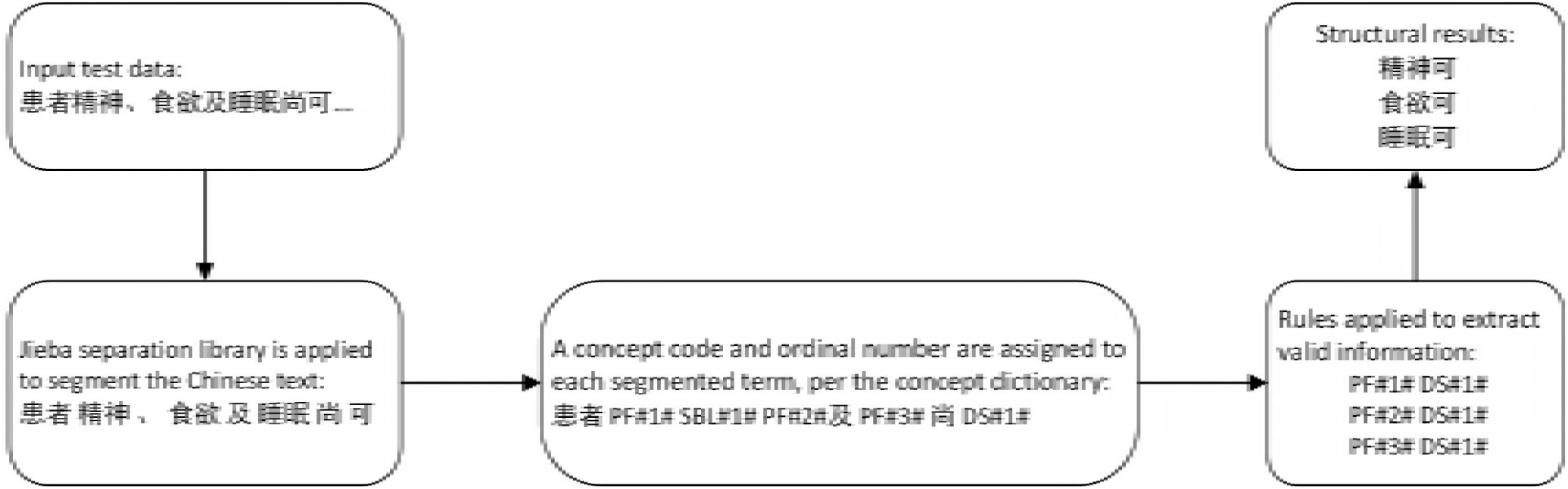
The extraction pipeline of the rule-based model.

#### Constructing the concept dictionary

The rule-based model requires the use of a Chinese medical word dictionary [33]. Thus, we constructed a lightweight Chinese medical word dictionary based on the EMR data set used in this experiment. Matching the key-value pairs of input text and dictionary concepts was crucial to the efficacy of the proposed approach. For example, the term “abdomen” corresponds to the concept “Body_Part,” while “cough” is recorded as a “Symptom.” The key-value matching process was repeated many times because the accuracy and the capacity of the concept dictionary directly affects the results of the subsequent rule-based structural extraction, especially the recall. Table 3 displays an excerpt of the concept dictionary constructed for this study.

**Table 3.** Excerpt of the concept dictionary with English translations.

| Segmented input (Chinese term) | English Translation | Concept |
| --- | --- | --- |
| 疼痛减轻 | Pain relief | @@Description |
| 尖索条影 | Tipstrip | @@Description |
| 节段性室壁运动 | Segmental Wall Motion | @@Symptom |
| 干湿啰音 | Dry and wet rales | @@Symptom |
| 腹型肥胖 | Abdominal obesity | @@Diagnosis |
| 未及 | Untouched | @@Diagnosis |
| 最大心率 | Maximum heart rate | @@Physiological_Function |
| 最小心率 | Minimum heart rate | @@Physiological_Function |
| 神志 | Spirit | @@Physiological_Function |
| 双侧颈动脉 | Bilateral carotid artery | @@Body_Part |
| 羟丁酸脱氢酶 | Hydroxybutyrate dehydrogenase | @@Lab_Test |
| 冠脉造影 | Coronary Angiography | @@Examination |
| 小便正常 | Urinary normal | @@Condition |

#### Rules

To establish the rules that would direct the structured extraction, we analyzed the EMR data set and mined rules based on commonly accepted medical transcription conventions. Thus, abstract sentences with similar attributes or content were classified into rules delivered as regular expressions [34]. Table 4 lists all the ten rules used by the proposed model. To identify additional meaningful patterns in the Chinese medical narratives, we created some more complex rules that consisted of two regular expressions.

**Table 4.**
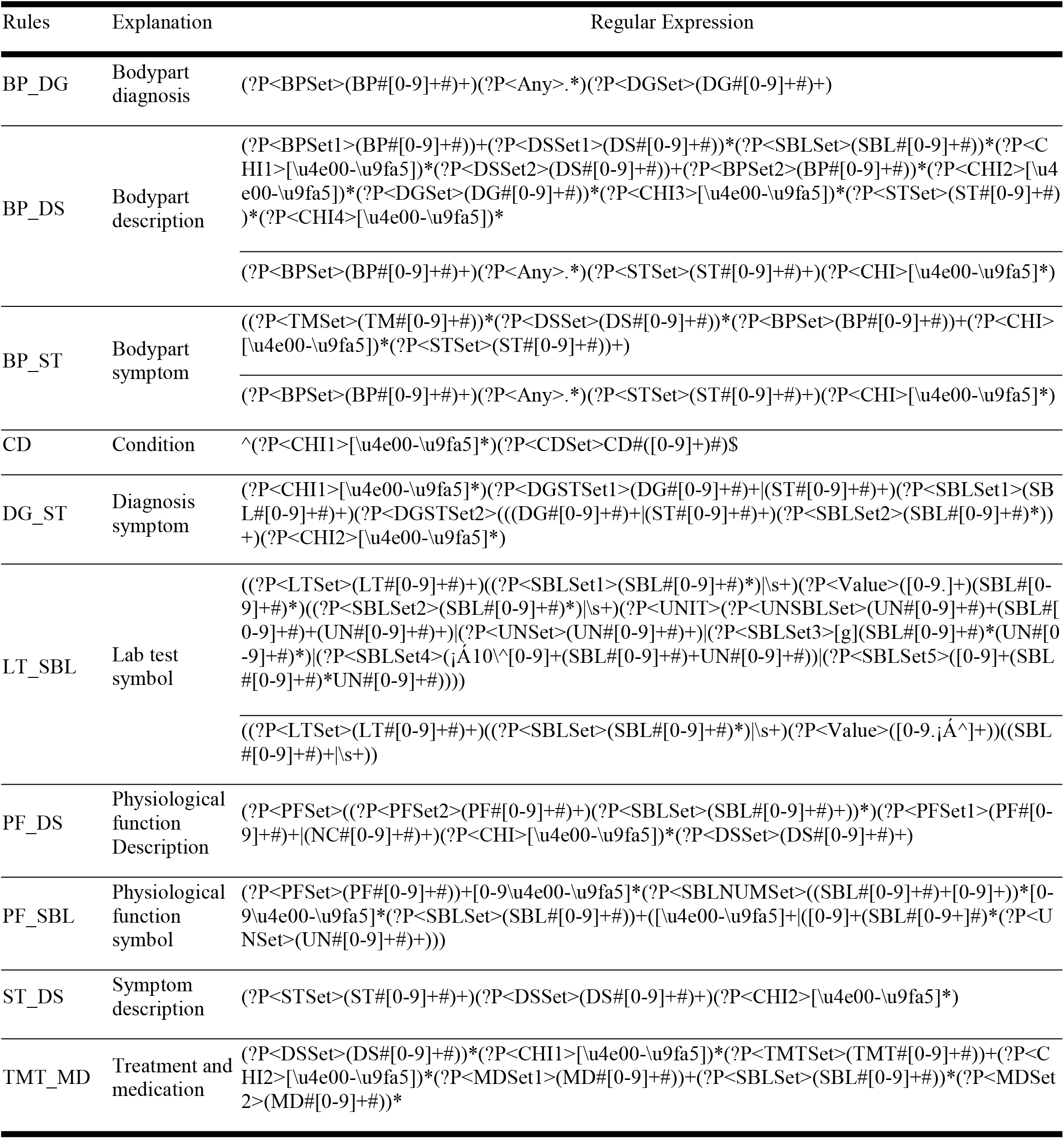
Rules used by the proposed rule-based model.

#### Revision of the extraction

The results of the structured extraction were constrained by the accuracy of the text segmentation. For example, if the segmentation was excessively fragmented, then the named entities of key-value pairs could be separated during segmentation, thus failing to recognize the appropriate conceptual pairing. Conversely, if the segmentation was too rough, the named entities could be embedded in larger segments that provide suboptimal specificity. Both situations would produce terms that could not be matched to regular expressions of the model rules and would therefore be excluded from the structured extraction implemented by enforcing these rules.

To mitigate omissions related to the granularity of the segmentation process, the extracted results were compared to a golden data set through multiple manual checks. During the debugging process, incorrect extraction results were evaluated to determine whether segmentation error was a factor. Regardless of the segmentation process, expected terms that were missing in the result set were added to an attached dictionary, and their frequency was noted. Frequency was an important consideration because higher frequency terms are easier to identify and properly extract to produce meaningful results. We note that the structured extraction results improved considerably after modifying the segmentation as described even though relatively few omissions were identified by the preceding revision process. The missing terms were added to the concept dictionary immediately after their discovery. As part of the process for incorporating the newly discovered terms, we tested whether the new terms could be correctly segmented, and adjusted the segmentation method as needed.

### The conditional random fields (CRF) model

#### Training model and test model

The conditional random field is a framework for building probabilistic models [35] widely used in natural language processing for word segmentation. The tagging set consists of EMRs processed by manual annotations. Four tags were assigned to the characters in a Chinese word: beginning (B), middle (M), end (E), or a single (S) word itself. While Chinese characters were described by BMES tags, punctuation and numbers were distinguished using the respective tags “PUNC” and “NUM.”

#### The pipeline of the conditional random fields (CRF) model

Fig 2 shows the extraction process of the CRF model. Note that the CRF extraction process that defines this model replaces the Jieba segmentation tool used by the preceding rules-based model. We first constructed the training model by manual annotating the EMR data set using the tags described in subsection 2.4.1. Then, the clauses and sentences segmented from the Chinese EMRs during preprocessing were supplied as input to the CRF model which executed word segmentation based on the assigned BMES tags. The subsequent structural extraction process resembled that of the rule-based model.

**Fig 2.**
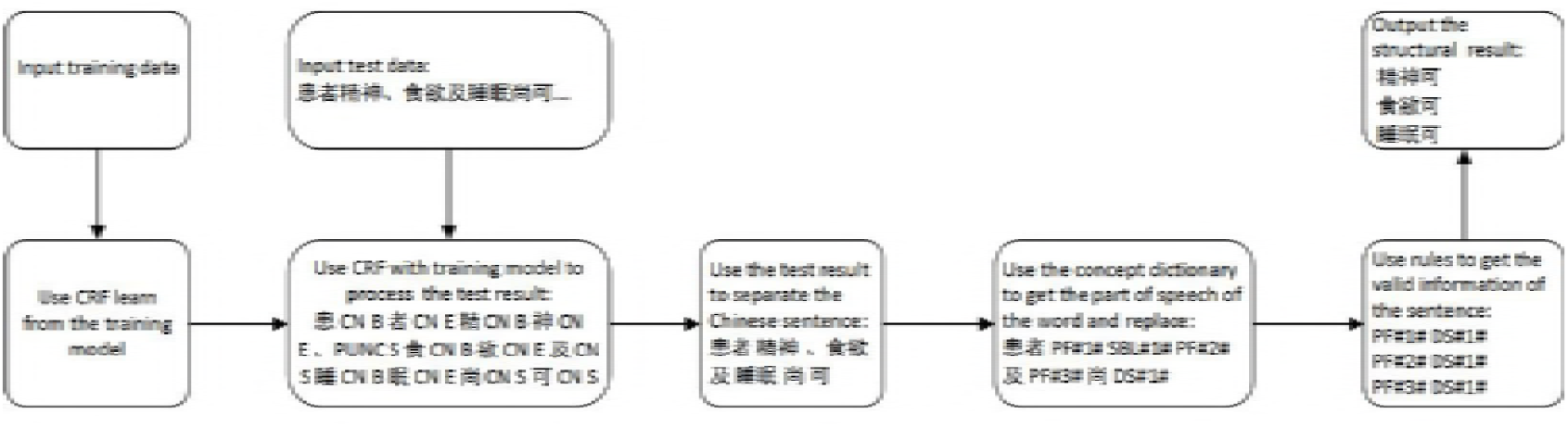
The extraction pipeline of the conditional random fields (CRF) model.

## Results

The performance of the proposed methods was evaluated using the standard three metrics, precision, recall, and F-score, derived from a confusion matrix. The input parameters of the confusion matrix are explained in Table 5. Table 6 provides the calculations used to obtain the statistical data and evaluation metrics presented in Table 7 and Table 8.

**Table 5.** Parameters for standard metrics.

| Parameters | Implication | Interpretation |
| --- | --- | --- |
| FN | False Negative | It is determined as a negative sample, but in fact a positive sample. |
| FP | False Positive | It is determined as a positive sample, but in fact a negative sample. |
| TN | True Negative | It is determined as a negative sample, and in fact a negative sample. |
| TP | True Positive | It is determined as a positive sample, and in fact a positive sample. |

**Table 6.** The calculation of some metrics in Table 7.

| Metrics | Calculation |
| --- | --- |
| Number of annotations | $TP + FN$ |
| Number of extractions | $TP + FP$ |
| Precision | $TP/(TP + FP)$ |
| Recall | $TP/(TP + FN)$ |
| F score | $2 \times Precision \times Recall / (Precision + Recall)$ |

**Table 7.** Statistical data measuring the performance of the rule-based model and the conditional random fields (CRF) model.

| Model | Number of annotations | Number of extractions | FP | TP | FN | Precision | Recall | F score |
| --- | --- | --- | --- | --- | --- | --- | --- | --- |
| Rule-based model | 1099 | **882** | 61 | 821 | 278 | 0.9308 | **0.7470** | 0.8289 |
| CRF-based model |  | 819 | 17 | 802 | 297 | **0.9792** | 0.7298 | **0.8363** |
| Rule-based model | 2041 | **1570** | 81 | 1489 | 552 | 0.9484 | **0.7295** | **0.8247** |
| CRF-based model |  | 1408 | 29 | 1379 | 662 | **0.9794** | 0.6756 | 0.7997 |
| Rule-based model | 3133 | **2412** | 102 | 2310 | 823 | 0.9577 | **0.7373** | **0.8332** |
| CRF-based model |  | 2151 | 46 | 2105 | 1128 | **0.9786** | 0.6719 | 0.7967 |
| Rule-based model | 3975 | **2972** | 123 | 2849 | 1126 | 0.9586 | **0.7167** | **0.8202** |
| CRF-based model |  | 2696 | 108 | 2588 | 1387 | **0.9599** | 0.6511 | 0.7759 |
| Rule-based model | 4626 | **3369** | 139 | 3230 | 1396 | 0.9587 | **0.6982** | **0.8080** |
| CRF-based model |  | 3189 | 113 | 3076 | 1550 | **0.9646** | 0.6649 | 0.7872 |

The bold data indicates the model with a better performance compared with each other.

**Table 8.**
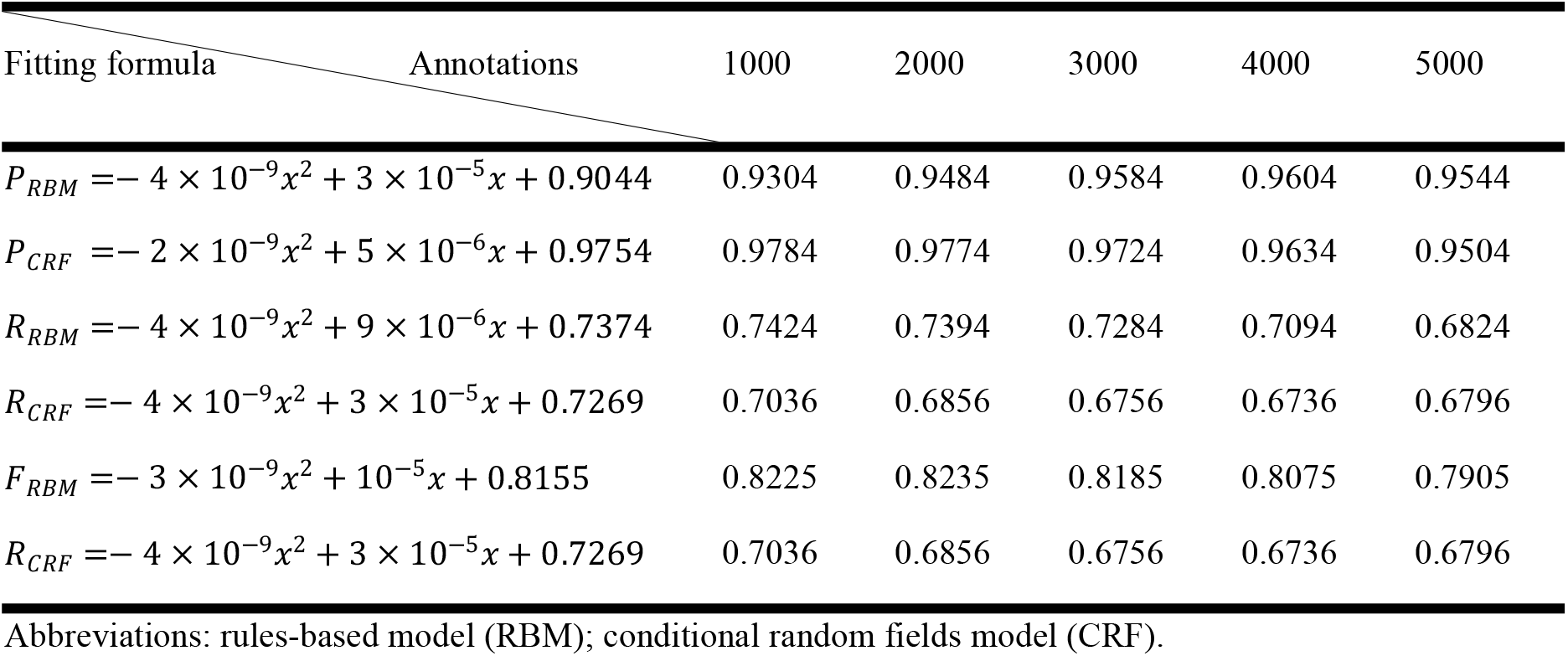
Statistical data of quadratic fitting curves.

Fig 3 and Fig 4 chart the variation and trendlines of precision, recall, and F-score as the annotations are increased using the rules-based model and the CRF model, respectively. The quadratic fitting curves are also plotted in the above Figs. According to the fitting formulas, calculating data is given in Table 8. And Fig 5 provides the comparison between two models.

**Fig 3.**
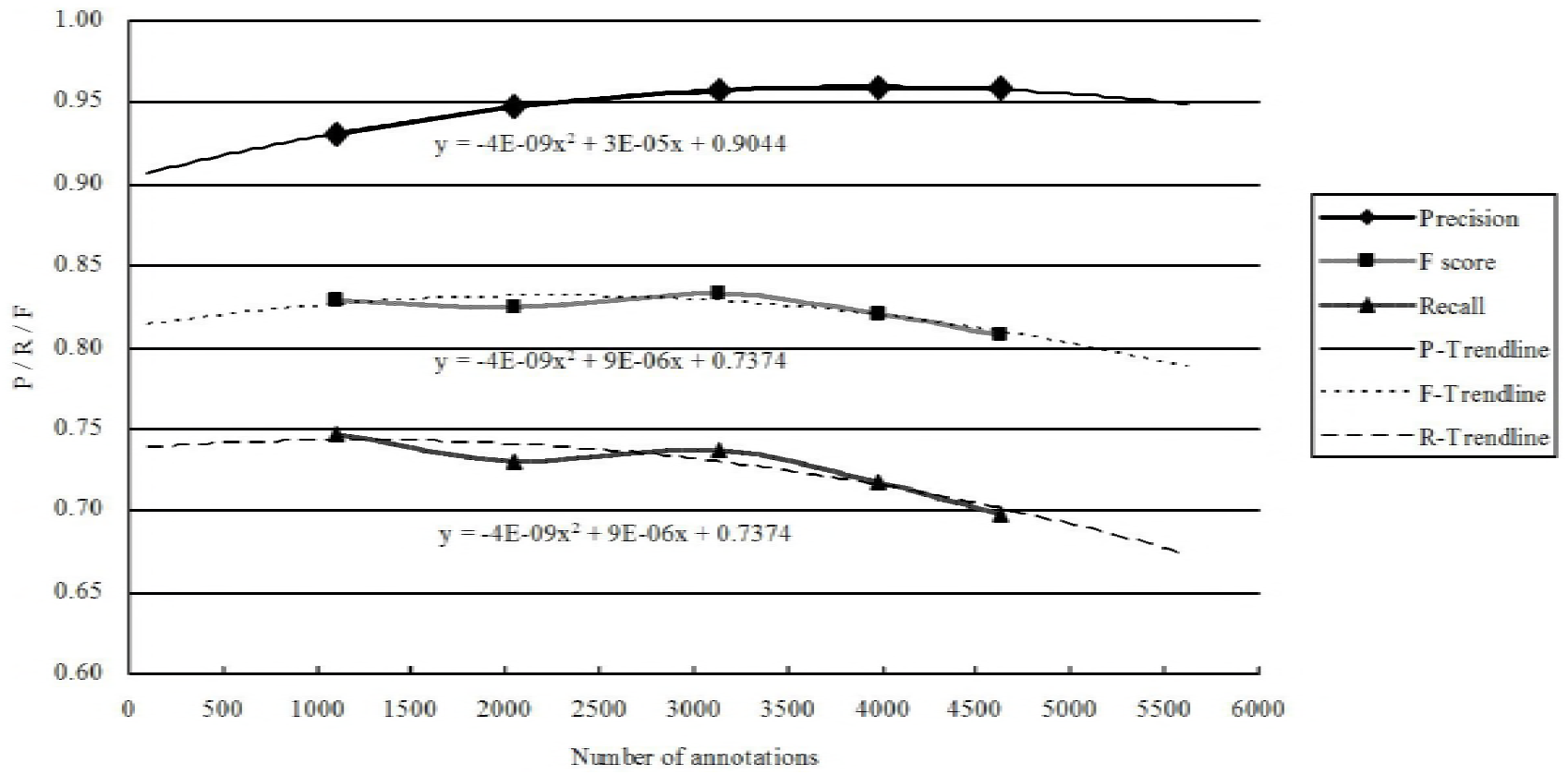
The variation tendency of extraction performance with the rule-based model.

**Fig 4.**
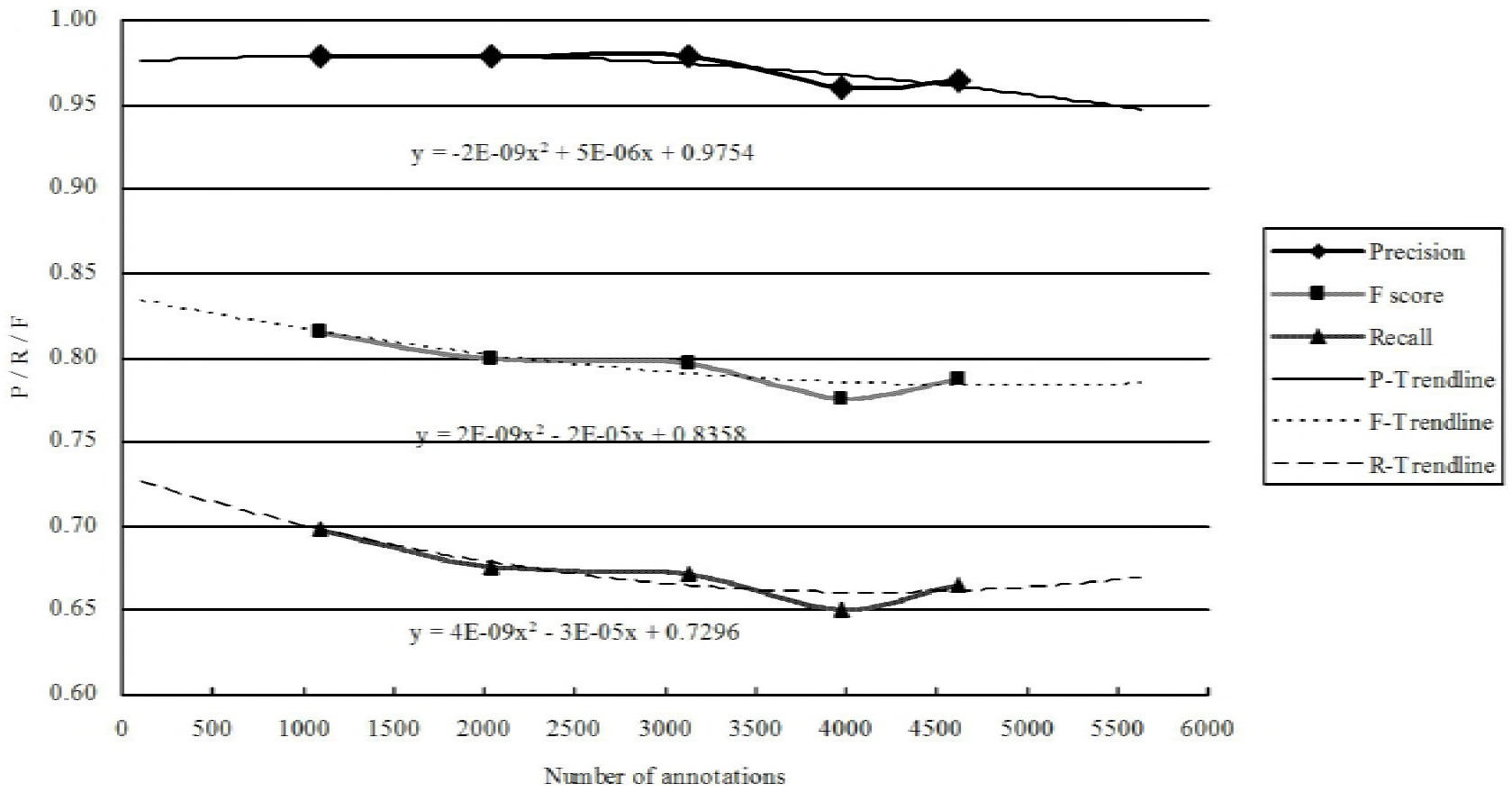
The variation tendency of extraction performance with the conditional random fields (CRF) model.

**Fig 5.**
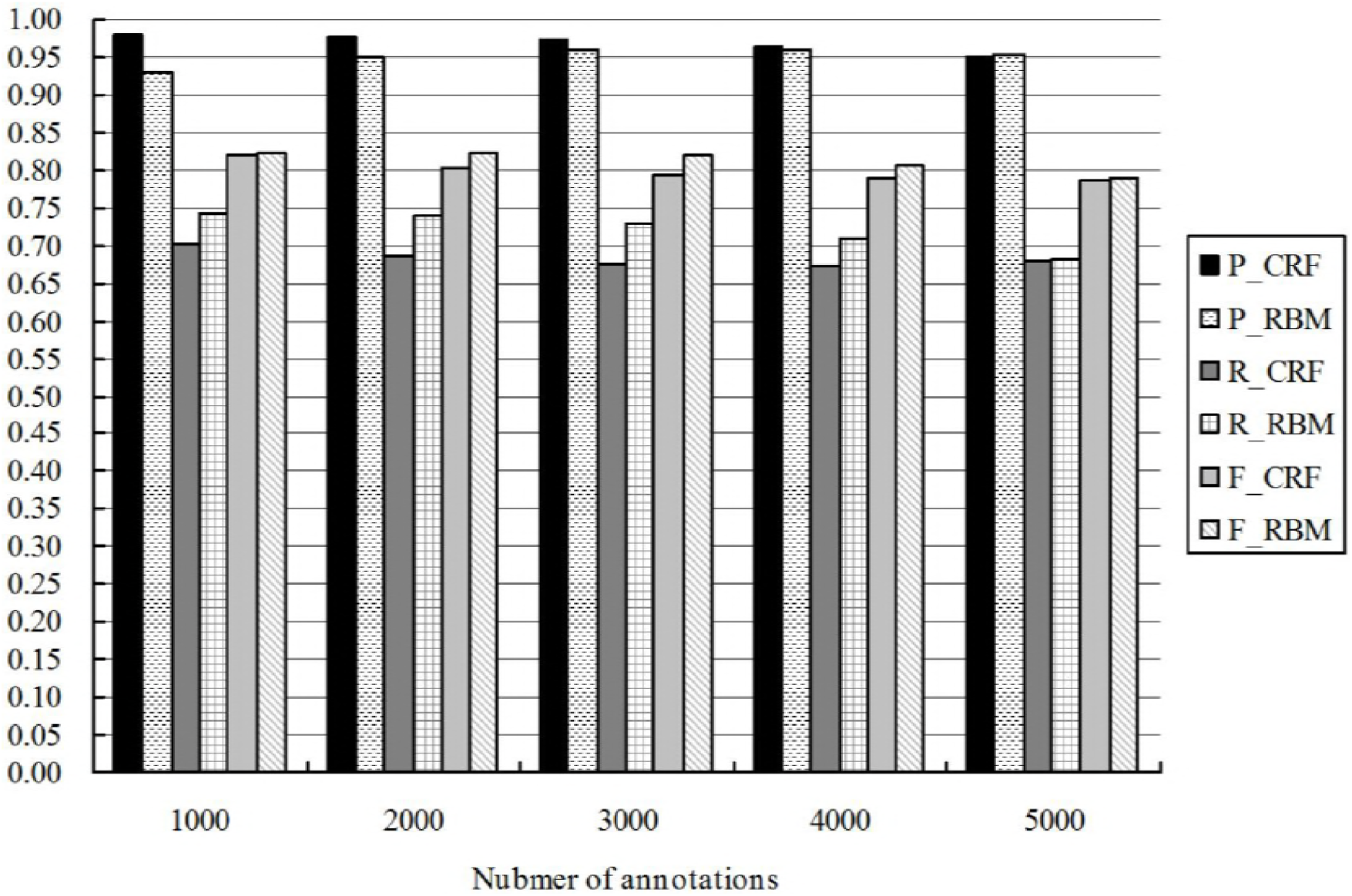
The comparison between fitting results of the conditional random fields (CRF) model and the rule-based model.

## Discussion

As the rule-based model is a white box method, the error statistics were used to determine whether a rule is acceptable or requires modification. Fig 6 graphs the error distribution per thousand units of annotated data, clearly indicating that the “Body parts-Description” (BP_DS) rule was the most flawed rule.

**Fig 6.**
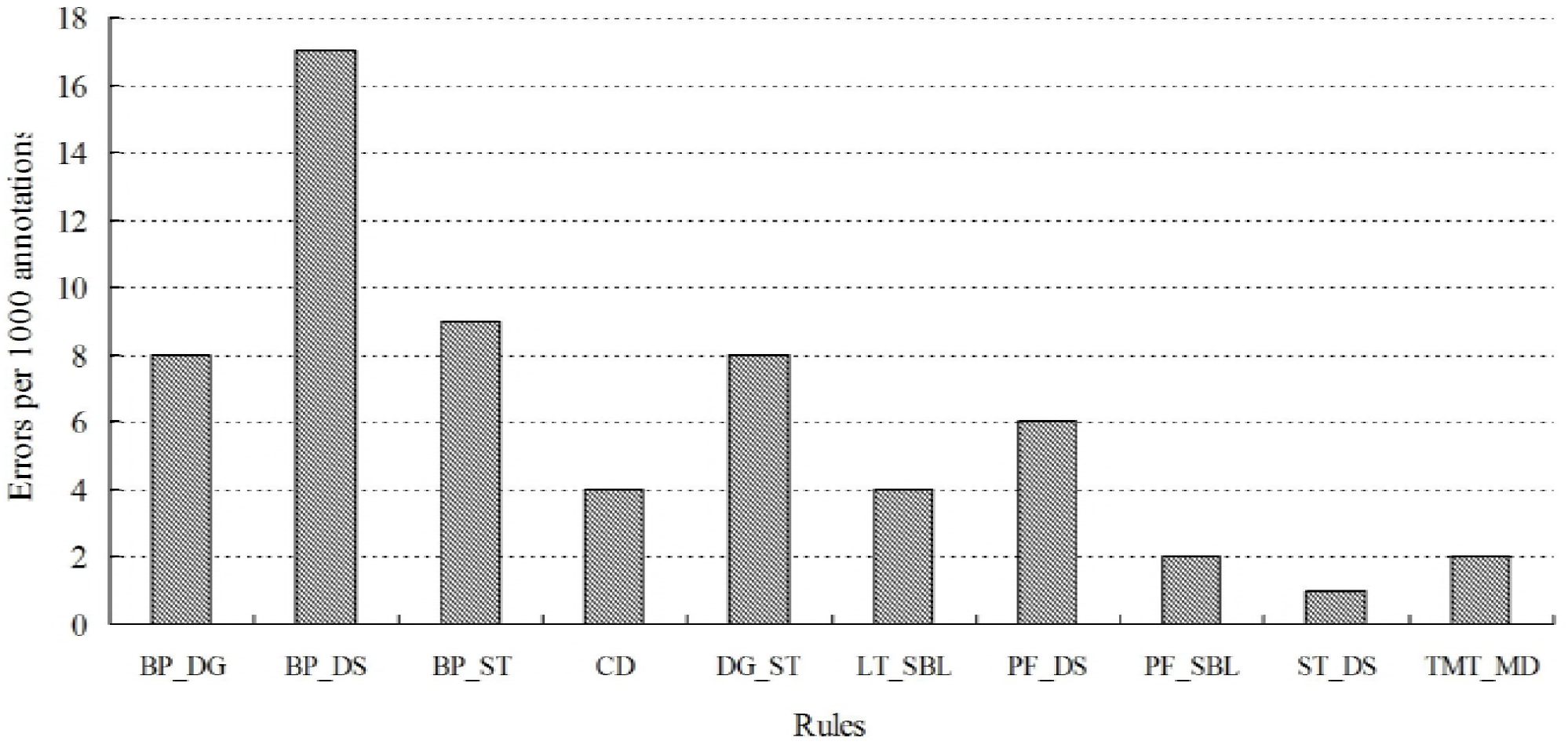
Errors caused by each rule per 1000 annotations. Abbreviations: BP_DG (Bodypart diagnosis); BP_DS (Bodypart description); BP_ST (Bodypart symptom); CD (Condition); DG_ST (Diagnosis symptom); LT_SBL (Lab test symbol); PF_DS (Physiological function Description); PF_SBL (Physiological function symbol); ST_DS (Symptom description); TMT_MD (Treatment and medication).

Table 9 analyzes an application of the BP_DS rule to show why the errors occurred. As observed in Table 9, faulty word segmentation caused the extraction error. In Chinese, “intervertebral” may also written as what translates to English as “intervertebral hole.” When the Chinese words were segmented, “intervertebral 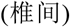” and “hole 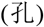” were separated. Thus, although the concept of “intervertebral 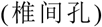 @@Body_Part” was in the concept dictionary, only the “narrow hole 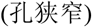” was extracted, resulting in the loss of semantics.

**Table 9.** Example 1 of error caused by BP DS rule.

|  | Extraction process in Chinese | Extraction process in English |
| --- | --- | --- |
| Original text | 椎间盘突出伴椎间孔狭窄 | Disc herniation with intervertebral foramen stenosis |
| Ideal extraction result | 椎间孔狭窄 | Intervertebral foramen stenosis |
| Actual extraction result<br>presented in XML format | `<RuleType Value="rule_BP_DS"> <Status Body_Part="孔" Condition="狭窄">True</Status> </RuleType>` | `<RuleType Value="rule_BP_DS"> <Status Body_Part="hole" Condition="stenosis">True</Status> </RuleType>` |
| Concept dictionary | 椎间孔@@Body_Part | intervertebral foramen@@Body_Part |
| Word segmentation | BP#7#:'孔','DS#8#:'狭窄' | BP#7#:'hole','DS#8#:'stenosis' |

Regarding the CRF model, our evaluation revealed that this model required a much larger training data set compared to the rule-based model. Despite using a training set of more than one million words, the training set did not include many common phrases and medical terms. Furthermore, the variety of word combinations and phrase sequences also exerted a strong influence on the correctness of the test set.

## Conclusions

This paper compared a rule-based model and a CRF model in extracting Chinese medical narratives. Among 4626 un-seen manually annotated records, the rules-based model achieved 95.87% precision, 69.82% recall, and an F-score of 80.80%, while the CRF model realized 95.99% precision, 65.11% recall, 77.59% F-score.

To summarize, for Chinese natural language texts with similar writing habits, such as progress notes and daily records, the narrative content is more easily abstracted into specific rules, to the advantage of the rule-based model. In contrast, although the recall of the CRF-based model is unremarkable, its accuracy is considerable. Increasing the size and quality of the training set would improve the performance of the CRF model. In the future work, we will consider an ensemble model that combines the merits of both the CRF model and the rule-based model. Furthermore, there is a great deal of unlabeled data, which could be used to extend the training set. Finally, it is also possible to apply a semi-supervised model to learn from both annotated data and unlabeled data to further improve the extraction results.

## Acknowledgements

This paper is supported by the Medical Informatics Lab at Zhejiang University, Hangzhou, Zhejiang, China.

## References

1. Chapman WW, Nadkarni PM, Hirschman L, D’avolio LW, Savova GK, Uzuner O. Overcoming barriers to NLP for clinical text: the role of shared tasks and the need for additional creative solutions. BMJ Group BMA House, Tavistock Square, London, WC1H 9JR; 2011.

2. Pestian JP, Brew C, Matykiewicz P, Hovermale DJ, Johnson N, Cohen KB, et al., editors. A shared task involving multi-label classification of clinical free text. Proceedings of the Workshop on BioNLP 2007: Biological, Translational, and Clinical Language Processing; 2007: Association for Computational Linguistics.

3. Uzuner Ö. Recognizing obesity and comorbidities in sparse data. Journal of the American Medical Informatics Association. 2009;16(4):561–70.

4. Uzuner Ö, Luo Y, Szolovits P. Evaluating the state-of-the-art in automatic de-identification. Journal of the American Medical Informatics Association. 2007;14(5):550–63.

5. Uzuner Ö, Solti I, Cadag E. Extracting medication information from clinical text. Journal of the American Medical Informatics Association. 2010;17(5):514–8.

6. Friedman C, Alderson P O, Austin J H M, et al. A general natural-language text processor for clinical radiology. Journal of the American Medical Informatics Association, 1994, 1(2): 161–174.

7. Sager N, Lyman M, Bucknall C, Nhan N, Tick LJ. Natural language processing and the representation of clinical data. Journal of the American Medical Informatics Association. 1994;1(2):142–60.

8. Hripcsak G, Friedman C, Alderson PO, DuMouchel W, Johnson SB, Clayton PD. Unlocking clinical data from narrative reports: a study of natural language processing. Annals of internal medicine. 1995;122(9):681–8.

9. Friedman C, Hripcsak G, DuMouchel W, Johnson SB, Clayton PD. Natural language processing in an operational clinical information system. Natural Language Engineering. 1995;1(1):83–108.

10. Haug PJ, Koehler S, Lau LM, Wang P, Rocha R, Huff SM, editors. Experience with a mixed semantic/syntactic parser. Proceedings of the Annual Symposium on Computer Application in Medical Care; 1995: American Medical Informatics Association.

11. Friedman C, Kra P, Yu H, Krauthammer M, Rzhetsky A, editors. GENIES: a natural-language processing system for the extraction of molecular pathways from journal articles. ISMB (supplement of bioinformatics); 2001.

12. Friedman C, Kra P, Yu H, et al. GENIES: a natural-language processing system for the extraction of molecular pathways from journal articles. ISMB (supplement of bioinformatics). 2001: 74–82.

13. Zou Q, Chu WW, Morioka C, Leazer GH, Kangarloo H, editors. IndexFinder: a method of extracting key concepts from clinical texts for indexing. AMIA Annual Symposium Proceedings; 2003: American Medical Informatics Association.

14. Zeng QT, Goryachev S, Weiss S, Sordo M, Murphy SN, Lazarus R. Extracting principal diagnosis, co-morbidity and smoking status for asthma research: evaluation of a natural language processing system. BMC medical informatics and decision making. 2006;6(1):30.

15. Wang X, Hripcsak G, Markatou M, Friedman C. Active computerized pharmacovigilance using natural language processing, statistics, and electronic health records: a feasibility study. Journal of the American Medical Informatics Association. 2009;16(3):328–37.

16. Murff HJ, FitzHenry F, Matheny ME, Gentry N, Kotter KL, Crimin K, et al. Automated identification of postoperative complications within an electronic medical record using natural language processing. Jama. 2011;306(8):848–55.

17. Grishman R, Sager N, Raze C, Bookchin B, editors. The linguistic string parser. Proceedings of the June 4–8, 1973, national computer conference and exposition; 1973: ACM.

18. Hirschman L, Grishman R, Sager N, editors. From text to structured information: automatic processing of medical reports. Proceedings of the June 7–10, 1976, national computer conference and exposition; 1976: ACM.

19. Sager N. Natural language information processing: A computer grammar of English and its applications. 1981.

20. Chi E, Lyman M, Sager N, Friedman C, Macleod C, editors. A database of computer-structured narrative: methods of computing complex relations. Proceedings of the Annual Symposium on Computer Application in Medical Care; 1985: American Medical Informatics Association.

21. Friedman C, Cimino JJ, Johnson SB, editors. A conceptual model for clinical radiology reports. Proceedings of the Annual Symposium on Computer Application in Medical Care; 1993: American Medical Informatics Association.

22. Pradhan S, Sun H, Ward W, Martin JH, Jurafsky D, editors. Parsing arguments of nominalizations in English and Chinese. Proceedings of HLT-NAACL 2004: Short Papers; 2004: Association for Computational Linguistics.

23. Qiu X, Zhang Q, Huang X, editors. Fudannlp: A toolkit for chinese natural language processing. Proceedings of the 51st Annual Meeting of the Association for Computational Linguistics: System Demonstrations; 2013.

24. Manning C, Surdeanu M, Bauer J, Finkel J, Bethard S, McClosky D, editors. The Stanford CoreNLP natural language processing toolkit. Proceedings of 52nd annual meeting of the association for computational linguistics: system demonstrations; 2014.

25. Huang C-R, Chen K-J, Chang L-L, editors. Segmentation standard for Chinese natural language processing. Proceedings of the 16th conference on Computational linguistics-Volume 2; 1996: Association for Computational Linguistics.

26. Che W, Li Z, Liu T, editors. Ltp: A chinese language technology platform. Proceedings of the 23rd International Conference on Computational Linguistics: Demonstrations; 2010: Association for Computational Linguistics.

27. Wang Y, Liu Y, Yu Z, Chen L, Jiang Y, editors. A preliminary work on symptom name recognition from free-text clinical records of traditional chinese medicine using conditional random fields and reasonable features. Proceedings of the 2012 Workshop on Biomedical Natural Language Processing; 2012: Association for Computational Linguistics.

28. Xu Y, Wang Y, Liu T, Liu J, Fan Y, Qian Y, et al. Joint segmentation and named entity recognition using dual decomposition in Chinese discharge summaries. Journal of the American Medical Informatics Association. 2013;21(e1):e84–e92.

29. Häyrinen K, Saranto K, Nykänen P. Definition, structure, content, use and impacts of electronic health records: a review of the research literature. International journal of medical informatics. 2008;77(5):291–304.

30. Abacha AB, Zweigenbaum P. Automatic extraction of semantic relations between medical entities: a rule-based approach. Journal of biomedical semantics. 2011;2(5):S4.

31. Patrick J, Li M. High accuracy information extraction of medication information from clinical notes: 2009 i2b2 medication extraction challenge. Journal of the American Medical Informatics Association. 2010;17(5):524–7.

32. Uzuner Ö, Solti I, Cadag E. Extracting medication information from clinical text. Journal of the American Medical Informatics Association. 2010;17(5):514–8.

33. Cimino JJ. From data to knowledge through concept-oriented terminologies: experience with the Medical Entities Dictionary. Journal of the American Medical Informatics Association. 2000;7(3):288–97.

34. Thompson K. Programming techniques: Regular expression search algorithm. Communications of the ACM. 1968; 11 (6):419–22.

35. Lafferty J, McCallum A, Pereira FC. Conditional random fields: Probabilistic models for segmenting and labeling sequence data. 2001.

